# Regulation of membrane scission in yeast endocytosis

**DOI:** 10.1101/2021.08.18.454332

**Authors:** Deepikaa Menon, Marko Kaksonen

## Abstract

During clathrin-mediated endocytosis, a flat plasma membrane is shaped into an invagination that undergoes scission to form a vesicle. In mammalian cells, the force that drives the transition from invagination to vesicle is primarily provided by the GTPase dynamin that acts in concert with crescent-shaped BAR domain proteins. In yeast cells, the mechanism of endocytic scission is unclear. The yeast BAR domain protein complex Rvs161/167 (Rvs) nevertheless plays an important role in this process: deletion of Rvs dramatically reduces scission efficiency. A mechanistic understanding of the influence of Rvs on scission however, remains incomplete. We used quantitative live-cell imaging and genetic manipulation to understand the recruitment and function of Rvs and other late-stage proteins at yeast endocytic sites. We found that arrival of Rvs at endocytic sites is timed by interaction of its BAR domain with specific membrane curvature. A second domain of Rvs167- the SH3 domain-affects localization efficiency of Rvs. We show that Myo3, one of the two type-I myosins in *Saccharomyces cerevisiae*, has a role in recruiting Rvs167 via the SH3 domain. Removal of the SH3 domain also affects assembly and disassembly of actin and impedes membrane invagination. Our results indicate that both BAR and SH3 domains are important for the role of Rvs as a regulator of scission. We tested other proteins implicated in vesicle formation in *Saccharomyces cerevisiae*, and found that neither synaptojanins nor dynamin contribute directly to membrane scission. We propose that recruitment of Rvs BAR domains delays scission and allows invaginations to grow by stabilizing them. We also propose that vesicle formation is dependent on the force exerted by the actin network component of the endocytic machinery.

## Introduction

Clathrin-mediated endocytosis is a process by which cargo molecules from the cell exterior are incorporated into a clathrin-coated vesicle that is then transported into the cell. Over 50 different proteins are involved in formation of an endocytic vesicle (***McMahon and Boucrot, 2011***; ***Kaksonen and Roux, 2018***). The molecular mechanisms mediating endocytic vesicle formation are largely conserved between yeast and mammals. Most yeast endocytic proteins have homologues in mammals. These proteins establish endocytic sites, recruit cargo, bend the membrane into an invagination, and finally separate the endocytic vesicle from the plasma membrane (***Kaksonen and Roux, 2018***). Actin filaments play a critical role in membrane-shaping in yeast. The filaments are nucleated and polymerize to form a branched actin network, which is required for bending the membrane into stereotypical tubular invaginations (***Kübler et al., 1993***; ***Kukulski et al., 2012***). How the final stage of membrane scission is effected in yeast remains unclear. In mammalian cells, the forces that drive the final transition from invagination to spherical vesicle are largely provided by the GTPase dynamin (***Grigliatti et al., 1973***; ***Takei et al., 1995***; ***Sweitzer and Hinshaw, 1998***; ***Ferguson et al., 2007***; ***Galli et al., 2017***). Dynamin interacts via its prolinerich domain with the SH3 domains of crescent-shaped N-BAR proteins like Endophilin and Amphiphysin (***Grabs et al., 1997***; ***Cestra et al., 1999***; ***Farsad et al., 2001***; ***Ferguson et al., 2009***; ***Meinecke et al., 2013***). Conformational changes of dynamin recruited to N-BAR molecules cause constriction of the underlying invaginated membrane, resulting in vesicle formation (***Shupliakov et al., 1997***; ***Zhang and Hinshaw, 2001***; ***Zhao et al., 2016***).

Three dynamin-like proteins: Dnm1, Mgm1, and Vps1, have been identified in yeast. Dnm1 and Mgm1 are involved in mitochondrial fusion and fission (***Cerveny et al., 2007***). The third, Vps1, gets its name from its essential role in vacuolar protein sorting in the secretory pathway (***Rothman et al., 1989***, ***1990***). It is also involved in fission and fusion of vacuoles and peroxisomes (***Hoepfner et al., 2001***; ***Peters et al., 2004***), and is required for regulation of endosome to Golgi trafficking (***Gurunathan et al., 2002***). In addition, Vps1 has been reported to localize at endocytic sites on the plasma membrane, interact with endocytic proteins like clathrin, and influence the lifetimes and recruitment of endocytic proteins (***Yu and Cai, 2004***; ***Nannapaneni et al., 2010***; ***Smaczynska-de Rooij et al., 2012***). Others however, have failed to observe Vps1 at endocytic sites (***Kishimoto et al., 2011***; ***Goud Gadila et al., 2017***), or observe a role in endocytic vesicle scission (***Nothwehr et al., 1995***; ***Kaksonen et al., 2005***). The role of Vps1 in clathrin-mediated endocytosis thus remains debated.

A confirmed component of the yeast endocytic scission mechanism is the heterodimeric complex formed by the N-BAR domain proteins Rvs161 and Rvs167 (***Munn et al., 1995***; ***D’Hondt et al., 2000***; ***Kaksonen et al., 2005***; ***Kishimoto et al., 2011***). The two Rvs proteins are homologues of the N-BAR proteins Amphiphysin and Endophilin in animals (***Friesen et al., 2006***; ***Youn et al., 2010***). Deletion of Rvs167 reduces scission efficiency by nearly 30% and reduces the lengths to which endocytic invaginations grow to nearly a third (***Kaksonen et al., 2005***; ***Kukulski et al., 2012***). In endocytic events that fail to undergo scission, the membrane first invaginates and then retracts back to the cell wall (***Kaksonen et al., 2005***). Rvs167 and Rvs161 proteins form a canonical N-BAR domain which forms a crescent-shaped structure (***Youn et al., 2010***). This curved structure is thought to be the key functional domain of the protein (***Sivadon et al., 1997***). N-BAR domains are able to form lattices that can bind membrane with their concave surfaces, and impose or sense curvature across dimensions larger than that of a single BAR domain (***Farsad et al., 2001***; ***Peter et al., 2004***; ***Youn et al., 2010***; ***Mim et al., 2012***; ***Zhao et al., 2013***). In addition to the N-BAR domain, Rvs167 has a Glycine-Proline-Alanine rich (GPA) region and a C-terminal SH3 domain (***Sivadon et al., 1997***). The GPA region is thought to act as a linker with no other known function, while loss of the SH3 domain affects daughter cell budding and actin cytoskeleton morphology (***Sivadon et al., 1997***). The Rvs complex can tubulate liposomes in vitro, indicating that the BAR domain can impose curvature on membranes (***Youn et al., 2010***). However, Rvs arrives at endocytic sites when membrane tubes are already formed (***Kukulski et al., 2012***; ***Picco et al., 2015***). Therefore curvature-sensing rather than curvature-generation is the likely role of the Rvs complex at endocytic sites. Rvs molecules arrive at endocytic sites a few seconds before scission, and disassemble rapidly at scission time (***Picco et al., 2015***), consistent with a role in vesicle scission. However, a mechanistic understanding of the role of Rvs in scission remains incomplete.

Synaptojanins are PI(4,5)P_2_ phosphatases that have been implicated in endocytic scission, as well as in intracellular signalling, and modulation of the actin cytoskeleton (***Singer-Krüger et al., 1998***). They interact with both dynamin and N-BAR proteins in mammalian cells (***McPherson et al., 1996***; ***Watanabe et al., 2018***). Disruption of these genes leads to accumulation of PI(4,5)P_2_ in cells (***Stolz et al., 1998b***). In synaptojain-disrupted mouse cells, coated endocytic vesicles cluster at the plasma membrane, demonstrating a role for lipid hydrolysis in vesicle uncoating (***Watanabe et al., 2018***). In yeast, removal of synaptojanin-like proteins affects rate of endocytosis, and induces aberrant behaviour of several endocytic proteins (***Singer-Krüger et al., 1998***; ***Sun et al., 2007***; ***Kishimoto et al., 2011***).

We aimed to identify roles and molecular mechanisms for proteins that have been implicated in endocytic vesicle scission in yeast: Vps1, synaptojanins, and the Rvs complex. We used quantitative live-cell imaging and genetic manipulation in *Saccharomyces cerevisiae* to test the roles of these proteins in endocytosis. We found evidence for a specific role in scission for the Rvs complex, but not for the other candidate proteins. Furthermore, we analyzed the molecular mechanisms of recruitment and the mode of action of Rvs in scission.

## Results

### Rvs167, but not Vps1, influences endocytic coat internalization

The role of the dynamin-like protein Vps1 in yeast endocytosis is unclear. Some studies have reported a role for Vps1 in endocytic vesicle scission (***Yu and Cai, 2004***; ***Nannapaneni et al., 2010***; ***Smaczynska-de Rooij et al., 2010***), while others have reported that Vps1 does not localize to endocytic sites or contribute to scission (***Kishimoto et al., 2011***; ***Goud Gadila et al., 2017***). Neither N- nor C-terminally tagged Vps1 co-localized with endocytic actin-binding protein Abp1 in our hands (data not shown), consistent with other work that observed localization only with the Golgi trafficking pathway (***Goud Gadila et al., 2017***). However, the question of whether or not Vps1 has a function at endocytic sites has been obfuscated by potential tagging-induced dysfunction of Vps1 molecules.

In mammalian cells, dynamin interacts with N-BAR proteins to cause vesicle scission (***Grabs et al., 1997***; ***Cestra et al., 1999***; ***Farsad et al., 2001***; ***Meinecke et al., 2013***). Although the association between yeast dynamin Vps1 and N-BAR protein Rvs is uncertain, Rvs is recruited to endocytic sites briefly before scission and influences scission efficiency (***Kaksonen et al., 2003***, ***2005***; ***Kukulski et al., 2012***; ***Picco et al., 2015***). In order to determine the roles of these proteins in endocytic scission, we analyzed the behaviour of other endocytic proteins in cells lacking Vps1 and Rvs167, and compared against wild-type (WT) cells (Fig.1A-F).

**Figure 1.**
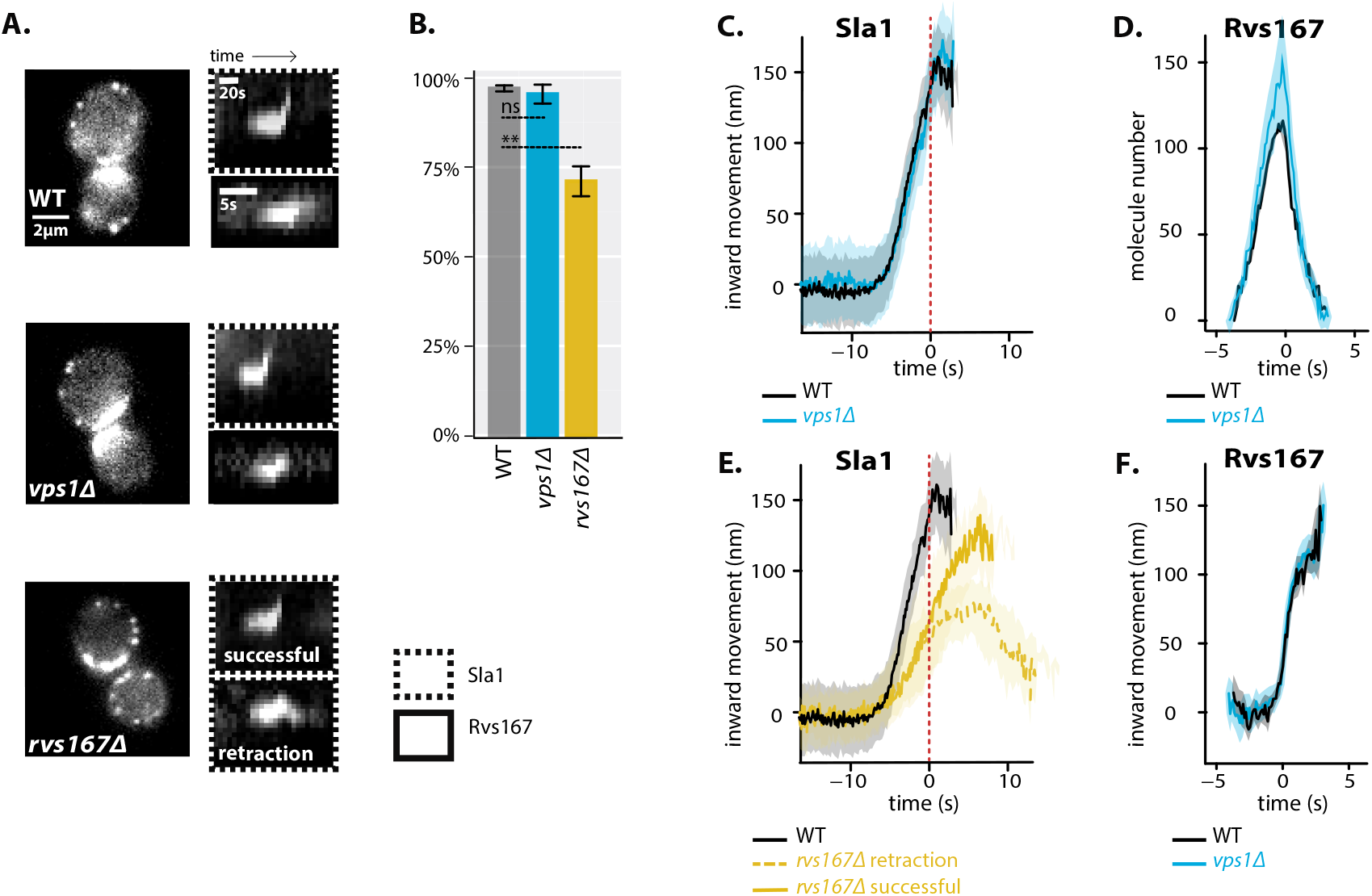
*vps1* Δ and *rvs167*Δ deletion. **A:** Left: Single frames from time-lapse movies of WT, *vps1*Δ, and *rvs167*Δ cells with endogenously tagged Sla1-eGFP. Right: Kymographs of Sla1-eGFP or Rvs167-eGFP in WT, *vps1*Δ, and *rvs167*Δ cells. **B:** Scission efficiency in WT, *vps1*Δ, and *rvs167*Δ cells. Error bars are standard deviation, p values from two-sided t-test, *= p ≤0.05, **= p≤0.01, ***=p≤0.001. **C:** Averaged centroid positions of Sla1-eGFP in WT and *vps1*Δ cells. **D:** Number of Rvs167 molecules in WT and *vps1*Δ cells. **E:** Averaged centroid positions of Sla1-eGFP in WT, and successful and retracted Sla1-eGFP positions in *rvs167*Δ cells. **F:** Averaged centroid positions of Rvs167-eGFP in WT and *vps1*Δ cells. All centroids were co-aligned with Abp1-mCherry so that time=0 s corresponds to Abp1 intensity maximum. On the y-axis, non-motile centroid position = 0 nm. Shading on plots show 95% confidence intervals. Dashed red lines indicate Abp1 intensity maxima in respective strains.

Vps1 gene deletion was confirmed by sequencing the gene locus. *vps1*∆ cells showed a previously reported slow growth phenotype at high temperatures (Fig.S1A) (***Rothman and Stevens, 1986***). In order to quantify invagination progression, coat protein Sla1 tagged at the C-terminus with eGFP was observed in yeast cells imaged at the equatorial plane (Fig.1A). Since membrane invagination progresses perpendicularly to the plane of the plasma membrane, proteins that move into the cytoplasm during invagination growth do so in the imaging plane. Upon actin polymerization, the endocytic coat moves into the cytoplasm along with the membrane as it invaginates (***Skruzny et al., 2012***). Movement of Sla1 thus acts as a proxy for the growth of the plasma membrane invagination. Membrane retraction, that is, inward movement and subsequent retraction of the invaginated membrane back towards the cell wall is a scission-specific phenotype (***Kaksonen et al., 2005***; ***Kishimoto et al., 2011***). Retraction rates were not significantly different in *vps1*∆ cells compared to the WT (Fig.1B).

To follow invaginations in more detail, centroid trajectories of 30-50 Sla1-eGFP patches in *vps1*∆ and WT cells were tracked and compared (Fig.1A-C). Centroids of Sla1 patches-each patch being an endocytic site-were tracked in time and averaged. This provided an averaged centroid that could be followed with high spatial and temporal precision (***Picco et al., 2015***). Sla1-eGFP was imaged simultaneously with Abp1-mCherry. Abp1 fluorescent intensity maximum in WT strains correlates with scission time and maximum movement of the Sla1 centroid (Fig.S1B) (***Picco et al., 2015***). The movement of the Sla1-eGFP centroid therefore corresponds to the growth of the endocytic membrane invagination and to the initial diffusive movement of the vesicle after scission (***Kukulski et al., 2012***; ***Picco et al., 2015***). The distance that Sla1 centroid moves thus gives an indication of invagination and scission steps of endocytosis. Any defects in scission are expected to chage the motility pattern of the Sla1 centroid.

The movement of Sla1-eGFP in WT cells was linear to about 150 nm (Fig.1C), consonant with maximum invagination lengths measured by correlative light and electron microscopy (CLEM) (***Kukulski et al., 2012***). Sla1 movement in *vps1*Δ cells, also co-aligned with Abp1-mCherry, was virtually identical to WT cells (Fig.1C). The total movement- and so the length of endocytic invagination - was similar to WT.

Quantitative imaging has shown that scission is simultaneous with a sharp jump of the Rvs167 centroid into the cytoplasm and a corresponding loss of fluorescent intensity (Fig.1 D,F, S1B) (***Kukulski et al., 2012***; ***Picco et al., 2015***). This jump is interpreted as loss of protein on the membrane tube at the time of scission, causing an apparent jump of the centroid to proteins that remain localized on the newly formed vesicle. Kymographs of Rvs167-GFP (Fig.1A), as well as Rvs167 centroid tracking (Fig.1F) in *vps1*Δ cells showed the same jump as in WT. We quantified the number of molecules of Rvs167 recruited to endocytic sites in *vps1*Δ cells (***Joglekar et al., 2006***; ***Picco et al., 2015***), and found that it was not significantly different from that recruited to WT cells (Fig.1D). We expect that a longer invagination is likely to recruit either more molecules of Rvs167 or the same number of molecules distributed along a longer invagination. Since we observe neither higher Rvs167 molecules numbers, nor larger invagination lengths, we conclude that the membrane tube is the same length as in WT. These data further suggest that the scission process is normal in *vps1*Δ cells.

We next studied invagination progression in cells lacking Rvs167. Since the Rvs complex is a dimer of Rvs161 and Rvs167 (***Boeke et al., 2014***), deletion of RVS167 gene effectively removes both proteins from endocytic sites (***Lombardi and Riezman, 2001***; ***Kaksonen et al., 2005***). We quantified the effect of *rvs167*Δ on membrane invagination (Fig.1A,B,E). When Sla1 was observed in *rvs167*Δ cells, nearly 30% of endocytic events displayed the beginning of movement away from the starting position- thus invagination formation- then retract back to starting position (Fig.1A,B). Retractions indicate failure of vesicle formation. Thus only 70% of Sla1 patches underwent apparently successful scission in *rvs167*Δ cells (Fig.1B). Similar scission rates have been measured in earlier studies (***Kaksonen et al., 2005***; ***Smaczynska-de Rooij et al., 2010***; ***Kishimoto et al., 2011***). We classified endocytic events into successful and retracting events and analyzed the average centroid movement in these two classes. Sla1 centroid movement in both successful and retracting endocytic events in *rvs167*Δ cells look similar to WT up to invagination length of about 50 nm (Fig.1E,S1E). In WT cells, Abp1 intensity begins to drop at scission time (Fig.S1B) (***Picco et al., 2015***). Abp1 intensity in *rvs167*Δ cells dropped after Sla1 centroid moved about 50 nm, suggesting that scission occurs in successful events at invagination lengths around 50 nm (Fig.S1D). That membrane scission occurs at shorter invagination lengths than in WT is corroborated by the smaller vesicles found using CLEM in *rvs167*Δ cells (***Kukulski et al., 2012***). CLEM has moreover shown that Rvs167 localizes to endocytic sites after the invaginations are about 50 nm long (***Kukulski et al., 2012***). Normal initial Sla1 movement in *rvs167*Δ indicates therefore that membrane invagination is unaffected till Rvs would normally arrive.

### Synaptojanins influence vesicle uncoating, but not scission dynamics

As Vps1 did not appear to influence membrane scission, we proceeded to test the potential role of synatojanins in scission (***Liu et al., 2009***). Apart from their role in vesicle uncoating, synaptojanins have been proposed to mediate scission with their PI(4,5)P_2_ hydrolysis activity (***Sun et al., 2007***; ***Toret et al., 2008***). In this model, BAR domains coat the invaginated tube, and preferential hydrolysis of PI(4,5)P_2_ at the invagination tip unprotected by BAR proteins generates line tension, eventually causing membrane scission. We reasoned that if the yeast synaptojanins are involved in scission, their deletion should alter the invagination dynamics visualized with Sla1-eGFP or Rvs167-eGFP. Three synaptojanin-like proteins have been identified in *S. cerevisiae*: Inp51, Inp52, and Inp53. Inp51-eGFP exhibits a diffuse cytoplasmic signal, Inp52-eGFP localizes to endocytic sites, and Inp53 localizes to patches within the cytoplasm (Fig.2A) (***Bensen et al., 2000***; ***Sun et al., 2007***). Since Inp52 can be observed at endocytic sites, we began with determining the spatial and temporal recruitment of Inp52 within the endocytic machinery. We tracked and aligned the averaged centroid of Inp52 spatially and temporally in relation to other endocytic proteins. In order to do this, we imaged Inp52-eGFP simultaneously with Abp1-mCherry. We also imaged Sla1-eGFP and Rvs167-eGFP together with Abp1-mCherry. Using Abp1 as the common reference frame, we were able to compare the arrival of the different proteins with respect to that of Abp1. We assigned as time =0 s, the peak fluorescent intensity of Abp1. In WT cells, this peak is concomitant with membrane scission, and also coincides with the peak Rvs167 fluorescent intensity (S1B) (***Picco et al., 2015***). On the y-axis, 0 nm indicates the non-motile position of the Sla1 centroid. Positions of the other centroids are spatially and temporally aligned to each other (Fig.2B). This analysis showed that Inp52 molecules arrived after Rvs167, and localized to the invagination tip. The localization and assembly dynamics of Inp52 are consistent with a role in the late stage of membrane invagination.

**Figure 2.**
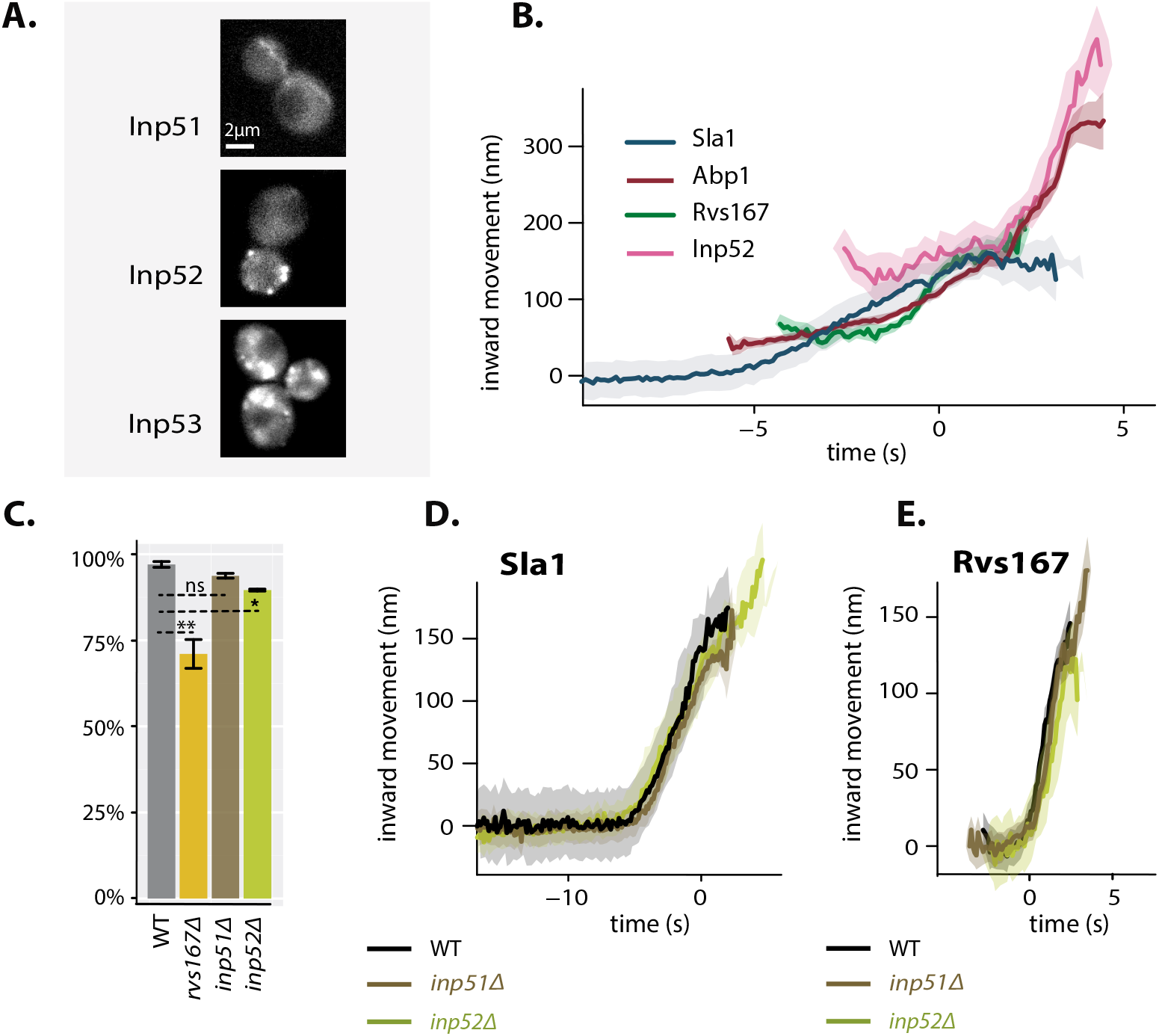
Synaptojanin-like proteins in yeast endocytosis. **A:** Single frames from time-lapse movies of cells with endogenously tagged Inp51-, Inp52-, and Inp53-eGFP. **B:**Inp52 centroid trajectory aligned in space and time to other endocytic proteins. **C:** Scission efficiency in WT, *rvs167*Δ, *inp51*Δ, and *inp52*Δ cells. Error bars are standard deviation, with p values from two-sided t-test, *= p≤0.05, **= p≤0.01, ***=p≤0.001. **D:** Averaged centroid positions of Sla1-eGFP in WT, *inp51*Δ, and *inp52*Δ cells. **E:** Averaged centroid positions of Rvs167-eGFP in WT, *inp51*Δ, and *inp52*Δ cells. D,E: On the y-axis non-motile Sla1 centroid position = 0 nm. All centroids were co-aligned with Abp1-mCherry so that time=0 s corresponds to Abp1 intensity maximum. Shading on plots represent 95% confidence interval.

Inp53 was not investigated further because it could not be detected at endocytic sites (Fig.2A), and is likely localized to the trans-Golgi network (***Bensen et al., 2000***). Although we were unable to observe localization of Inp51 at the plasma membrane (Fig.2A), deletion of Inp51 has been shown to exacerbate the effect of *inp52*Δ on endocytosis (***Singer-Krüger et al., 1998***), so both Inp51 and Inp52 were tested as potential scission regulators.

*inp51*Δ*inp52*Δ cells have dramatic morphological and growth defects, defects in vacuole morphology and budding polarity (***Singer-Krüger et al., 1998***; ***Stolz et al., 1998a***). These cells also have drastically altered PI(4,5)P_2_ levels (***Stolz et al., 1998b***), which likely affect the assembly, disassembly, and function of many PI(4,5)P_2_-binding endocytic proteins. The double mutation reportedly causes aberrations in endocytic coat, myosin, and actin network behaviour (***Sun et al., 2007***). Coat proteins Sla1, Sla2 and Ent1 have elongated lifetimes at endocytic sites, as do type I myosin Myo5, and Rvs167. Time taken for Abp1 assembly and disassembly is more than doubled (***Sun et al., 2007***). That multiple endocytic phases, including scission, are affected in the double mutation makes it difficult to demonstrate a direct role in scission. Patches of Rvs167-eGFP tracked in these cells persist instead of disassembling immediately after inward movement, leading to aggregation of fluorescent patches inside the cytoplasm (Fig.S2B).

We cannot, by the methods used here, distinguish between scission and other defects. We reasoned that a quantitative analysis of single mutants was therefore better suited to reveal a scission-specific function for synaptojanins, without perturbing overall PI(4,5)P_2_ homeostasis.

Dynamics of Sla1-eGFP and Rvs167-eGFP in *inp51*Δ and *inp52*Δ cells were compared against the WT (Fig.2C-E). Scission efficiency did not significantly decrease in *inp51*Δ compared to the WT, but showed a slight decrease in *inp52*Δ cells (Fig.2C). The movement of Sla1 and Rvs167 centroids in successful endocytic events in *inp51*Δ were virtually the same as in WT (Fig.2 D,E), while Rvs167 assembly and disassembly took longer (Fig.S2A). Rvs167 signal in *inp51*Δ cells persisted longer compared to the WT (Fig.2E), likely because of a delay in Rvs167 disassembly from the newly formed vesicle. In *inp52*Δ cells, Sla1 centroid movement had the same magnitude and rate as in WT, but Sla1-eGFP signal was persistent after inward movement (Fig.2D). Sla1 assembly and disassembly were aberrant in *inp52*Δ cells compared to WT (Fig.S2A). These data are consistent with synaptojanin involvement in assembly and disassembly of coat and scission proteins at endocytic sites (***Toret et al., 2008***). However, because the centroid movements of Sla1 and Rvs167 are unaltered, synaptojanins may not have a direct or major role in membrane scission.

### Rvs BAR domains recognize membrane curvature in vivo

So far Rvs167 and Rvs161 remain the proteins that have the most significant influence on scission efficiency. Recruitment to and interaction of the Rvs complex at endocytic sites was thus investigated further. The Rvs complex can tubulate liposomes in vitro, likely via the BAR domain (***Youn et al., 2010***). Interaction of the BAR domain with membrane curvature in vivo has however not been tested. The Rvs167-SH3 domain can interact with proteins associated with actin patches such as Abp1, Las17, Myo3, Myo5 and Vrp1, but the role of these interactions in vivo is not known (***Lila and Drubin, 1997***; ***Colwill et al., 1999***; ***Madania et al., 1999***; ***Liu et al., 2009***). We first tested the BAR-membrane interaction by deleting the SH3 domain to remove its contribution (Fig.3A). We then observed localization of Rvs167-sh3Δ in *sla2*Δ cells, which do not have membrane curvature at endocytic sites (***Picco et al., 2018***). Sla2 acts as the molecular linker between forces exerted by the actin network and the plasma membrane (***Skruzny et al., 2012***). *sla2*Δ cells therefore contain a polymerizing actin network at endocytic patches, but the membrane has no curvature, and endocytosis fails (***Skruzny et al., 2012***; ***Picco et al., 2018***). Colocalization of endogenously tagged full-length Rvs167-eGFP and Rvs167-sh3Δ-eGFP with Abp1-mCherry in WT and *sla2*Δ cells were compared (Fig.3B). We thus tested whether Rvs BAR domain could be recruited to the endocytic sites in *sla2*Δ cells, independent of membrane curvature. In *sla2*Δ cells, the full-length Rvs167 co-localized with Abp1-mCherry indicating that it was recruited to endocytic sites without membrane curvature (Fig.3B, “*sla2*Δ”). Rvs167-sh3Δ did not appear at the plasma membrane except for rare transient patches. Therefore, Rvs167-sh3Δ, that is, the BAR domain alone, is not recruited to endocytic sites in the absence of curvature in *sla2*Δ cells. Localization of the full-length protein in *sla2*Δ cells is therefore likely via SH3 domain interaction.

**Figure 3.**
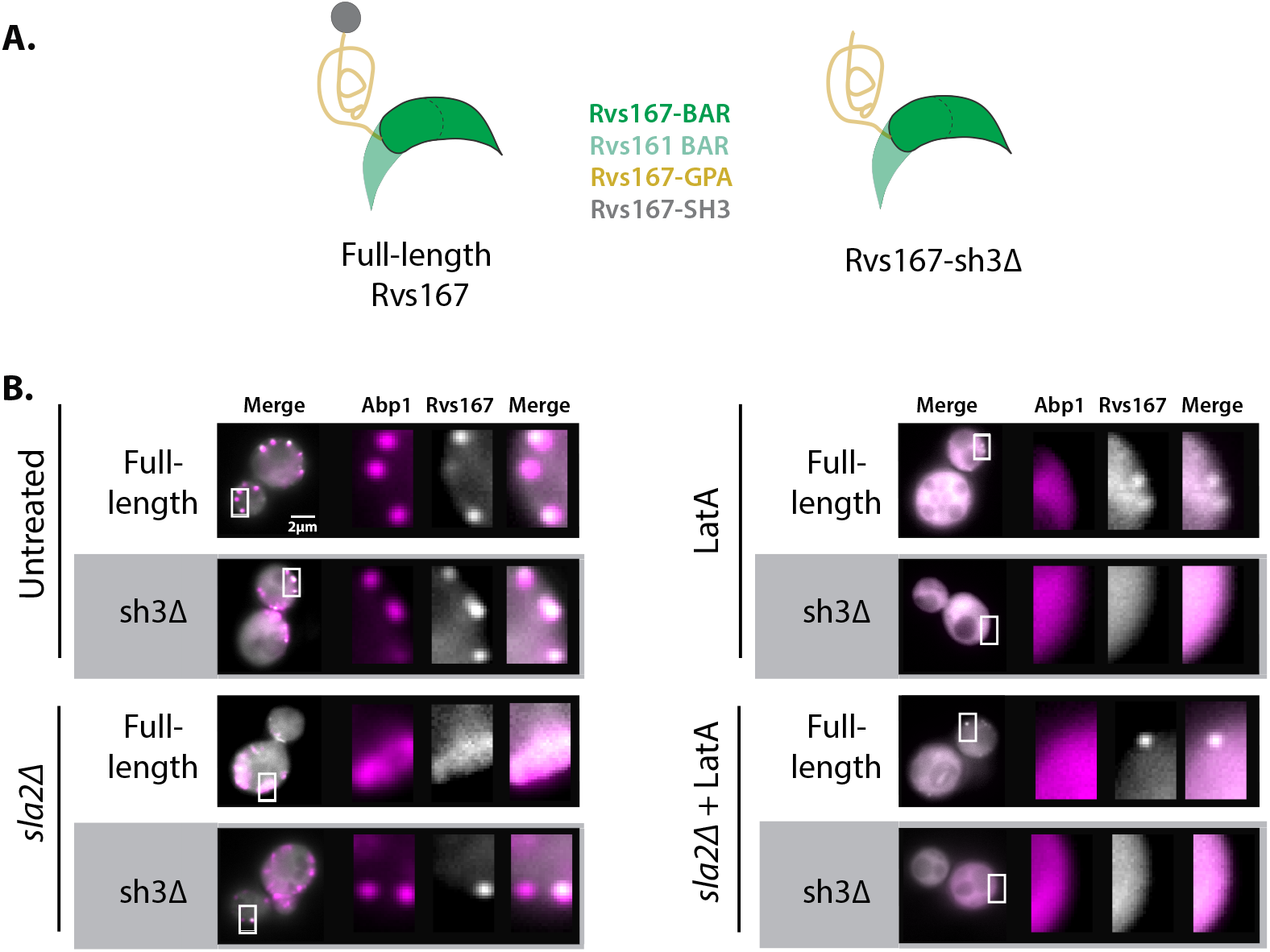
Localization of Rvs167 BAR domain. **A:** Schematic of Rvs protein complex with and without the SH3 domain. **B:** Localization of full-length Rvs167 and Rvs167-sh3Δ in WT, *sla2*Δ, LatA treated, and LatA treated *sla2*Δ cells.

### Rvs SH3 domains have an actin and curvature-independent localisation

We wanted to distinguish between Rvs association with membrane curvature and actin. Latrunculin A (LatA) inhibits actin polymerization, and therefore the assembly of actin and actin-related proteins at endocytic sites. *sla2*Δ as well as LatA remove membrane curvature, but *sla2*Δ retains actin patches at endocytic sites (***Kukulski et al., 2012***; ***Picco et al., 2018***). To study the interaction of Rvs at endocytic sites without actin proteins, we observed the localization of Rvs167 and Rvs167-sh3Δ in LatA treated WT cells (Fig.3B, “LatA”). We also observed full-length and mutant protein in *sla2*Δ cells treated with LatA, so that we could be sure that we removed any capacity for membrane curvature, as well as actin related proteins (Fig.3B, “*sla2*Δ + LatA”). Full-length Rvs167 localized transiently at the plasma membrane in WT cells treated with LatA as well as in *sla2*Δ cells treated with LatA (Fig.3B). Rvs167-sh3Δ did not localize to the plasma membrane in either case. Thus, localization of full-length Rvs167 in the presence of LatA in WT and *sla2*Δ cells is due to the SH3 domain. This also indicates that the SH3 domain is able to recruit Rvs molecules to the plasma membrane in an actin- and curvature-independent manner.

### Rvs167 SH3 domains are likely recruited by Myo3

Type I myosins Myo3 and Myo5, and yeast verprolin Vrp1 have known genetic or physical interactions with the Rvs167 SH3 domain (***Lila and Drubin, 1997***; ***Colwill et al., 1999***; ***Madania et al., 1999***; ***Liu et al., 2009***). We tested the possible role of these proteins in the SH3-dependent localization of Rvs167 in cells with the gene for one of these proteins deleted, and treated with LatA (Fig.4). By using LatA we expected to reproduce the situation in which the interaction between the BAR domain and curved membrane is removed. Then, if we lost SH3 interaction because we removed the protein with which it interacts, we would lose localization of Rvs167 completely. Neither Vrp1 nor Myo5 deletion in combination with LatA treatment removed the localization of Rvs167: 88% and 87% cells respectively still showed Rvs167 localization, similar to WT localization. Deletion of Myo3 with LatA treatment reduced localization of Rvs167 (43.4% of cells with localization), indicating that SH3 domains interact at endocytic sites primarily with Myo3.

**Figure 4.**
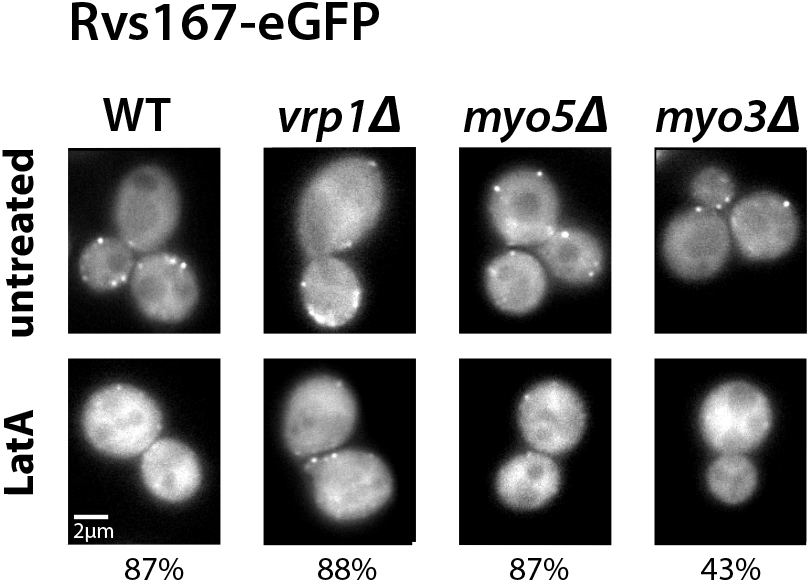
Localization of Rvs167 in the absence of membrane curvature. Single frames from time-lapse movies showing Rvs167-eGFP localization in untreated and LatA treated WT, *vrp1*Δ, *myo5*Δ, and *myo3*Δ cells. Percentages indicate number of LatA treated cells in which Rvs167-eGFP is localized at the plasma membrane.

### Deletion of Rvs167 SH3 domain affects coat and actin dynamics

Since the Rvs167-SH3 domain had an influence on the recruitment of the Rvs complex to endocytic sites, we wondered if the domain affects not just recruitment of the protein, but also invagination progression. We compared dynamics of coat, actin, and scission markers in WT and *rvs167-sh3*Δ cells (Fig.5).

**Figure 5.**
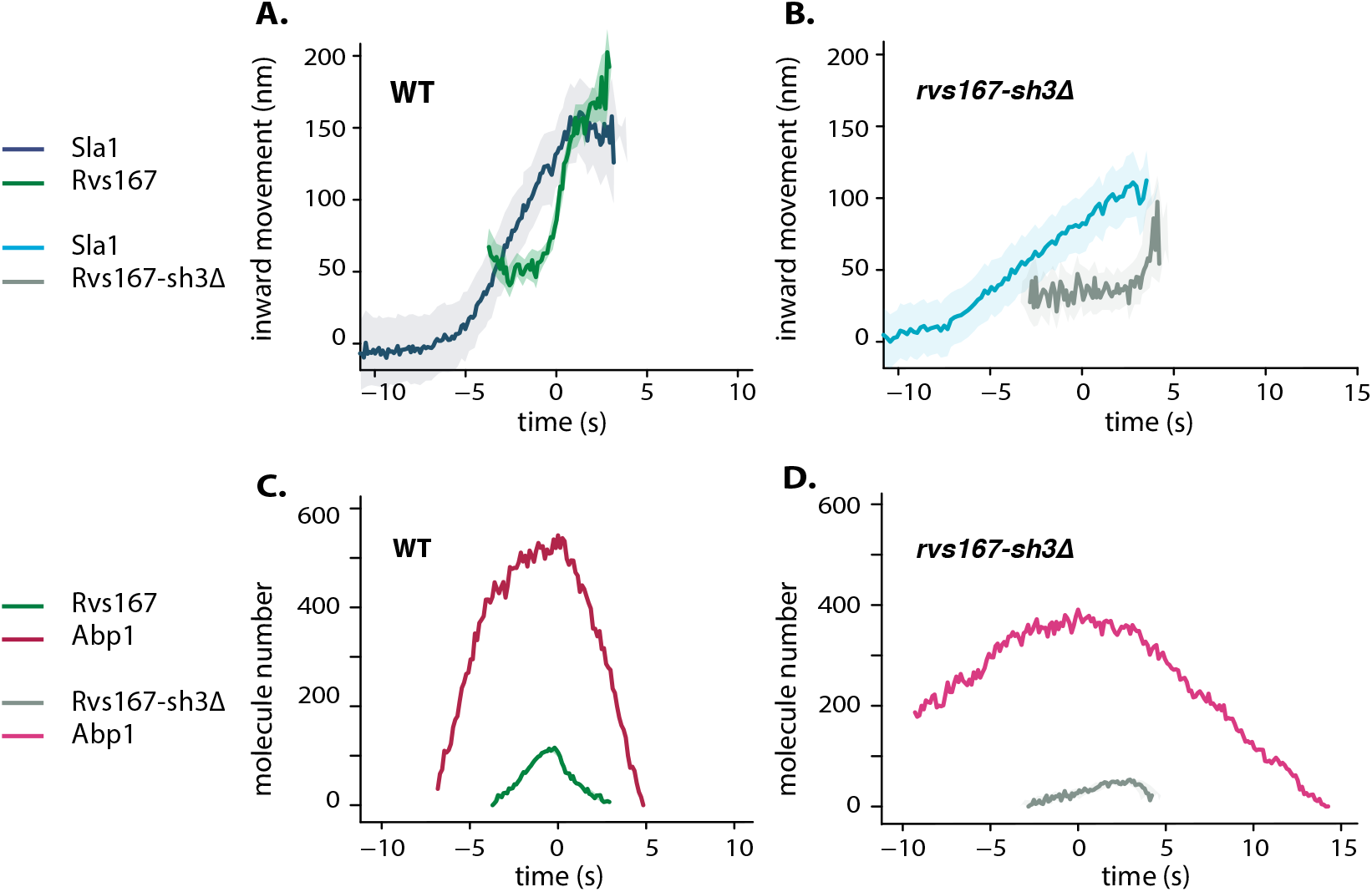
Endocytic dynamics in WT and *rvs167-sh3* Δ cells. **A,B:** Averaged centroid positions aligned in x-axis so that time=0 s corresponds to Abp1 intensity maximum in the respective strains. Centroids were aligned in y-axis so that non-motile Sla1 position is at y=0 nm, and Rvs167 and Rvs167-sh3Δ positions are determined with respect to Sla1 centroids. **C,D:** Numbers of molecules in WT and *rvs167-sh3*Δ cells, aligned so that time=0 s is the Abp1 intensity maximum in the corresponding strains. Shading represents 95% confidence interval.

The Sla1 centroid position in *rvs167-sh3*Δ cells at scission time is about 70 nm compared to 150 nm in WT, and this movement takes 8 s compared to 5 s in WT (Fig.5A,B). The total movement of Rvs167-sh3Δ centroid is half that of full-length Rvs167 (Fig.5A,B). Reduced movements of both Sla1 and Rvs167-sh3Δ centroids in *rvs167-sh3*Δ cells are consistent with the formation of shorter endocytic invaginations in these cells.

We observed that Rvs167-sh3Δ recruitment begins at nearly the peak of Abp1 recruitment in *rvs167-sh3*Δ cells, while in WT, full-length Rvs167 is recruited halfway into Abp1 recruitment (Fig.5 C,D). Rvs167-sh3Δ accumulation however, began when Abp1 molecule number in the mutant was the same as in WT (300 copies, Fig.5C,D). Both Rvs167 and Rvs167-sh3Δ molecules arrived at endocytic sites when the Sla1 centroid was 30-50 nm away from its starting position, so the endocytic coat has moved a certain amount when both WT and mutant forms of Rvs start to be recruited. That Rvs167-sh3Δ recruitment begins at a certain length of the invagination suggests that the Rvs complex is recruited to a specific geometry of membrane invagination. Rvs167-sh3Δ accumulation may be delayed because invaginations in these cells take longer to acquire this geometry.

We think that Rvs molecules in WT cells likely arrive below our detection threshold, and that the arrival of the molecules supports invagination growth. As the invagination grows, Rvs continues to accumulate on the invagination tubes, and molecule numbers are eventually large enough to be detected. In support of this, CLEM has shown that when Rvs167 molecules are detected at endocytic sites, the invaginations are about 50 nm long, shortly after the membrane is already tubular (***Kukulski et al., 2012***; ***Picco et al., 2015***). Since Rvs167-sh3Δ molecules accumulate slower than full-length protein, their support to membrane growth is less effective, and the invagination grows slower. Abp1 accumulation correlates with invagination growth, so slower invagination growth accumulates Abp1 slower. Recruitment of Rvs167-sh3Δ was significantly reduced compared to Rvs167 (Fig.5C,D), although cytoplasmic concentration of both were similar (Fig.S4). Recruitment therefore is unlikely to be limited by expression of the mutant protein. Abp1 disassembly time was increased to 15 s in *rvs167-sh3*Δ cells compared to 5 s in WT, and total number of Abp1 molecules recruited was reduced from nearly 600 to 400, 60% of WT recruitment (Fig.5C,D). Recruitment and disassembly defects of Abp1 indicate disruption of actin network dynamics in *rvs167-sh3*Δ cells.

In WT cells, the number of Rvs167 and Abp1 molecules peak at the same time. Thus, the actin network begins disassembling as soon as scission occurs (Fig.5C). However, in *rvs167-sh3*Δ cells, the numbers of Rvs167 and Abp1 molecules peaked asynchronously, with Rvs167 peaking later. This observation suggests that there is a feedback mechanism between the actin network and membrane scission, and that this feedback is also disrupted in *rvs167-sh3*Δ cells.

### Increased BAR domain recruitment corresponds to increased membrane movement

Reduced Sla1 movement was observed in both *rvs167*Δ (Fig.1) and *rvs167-sh3*Δ (Fig.5) cells, in which about half the WT number of Rvs167 molecules are recruited (Fig.5). This suggests that increased Sla1 movement correlates with increased recruitment of Rvs167. We wondered if Sla1 movement would scale with amount of Rvs recruited to endocytic sites. This could suggest that recruitment of Rvs BAR domains scaffolds the membrane invagination and protects it against membrane scission (***Boucrot et al., 2012***; ***Dmitrieff and Nédélec, 2015***). We titrated the amount of Rvs expressed in cells by duplicating the open reading frame of RVS167 and RVS161 genes (***Huber et al., 2014***). We also generated a strain in which the *rvs167-sh3*Δ gene was duplicated. We thus obtained cells containing either 2x copies of both RVS genes (2xRVS), 1x copy of the RVS genes (1xRVS, ie WT), 2x copies of *rvs167-sh3*Δ (2xBAR), or 1x copy of *rvs167-sh3*Δ (1xBAR) (Fig.6A-D). In the 2xBAR strain, RVS161 was not duplicated. This is because we measured the number of molecules of mutant Rvs167 recruited in the 2xBAR strain with and without RVS161 duplicated, and found that they were the same, suggesting that Rvs161 protein expression is not limiting for assembly of the Rvs complex in this strain (data not shown). So we used the genetically simpler strain, without the RVS161 duplication.

**Figure 6.**
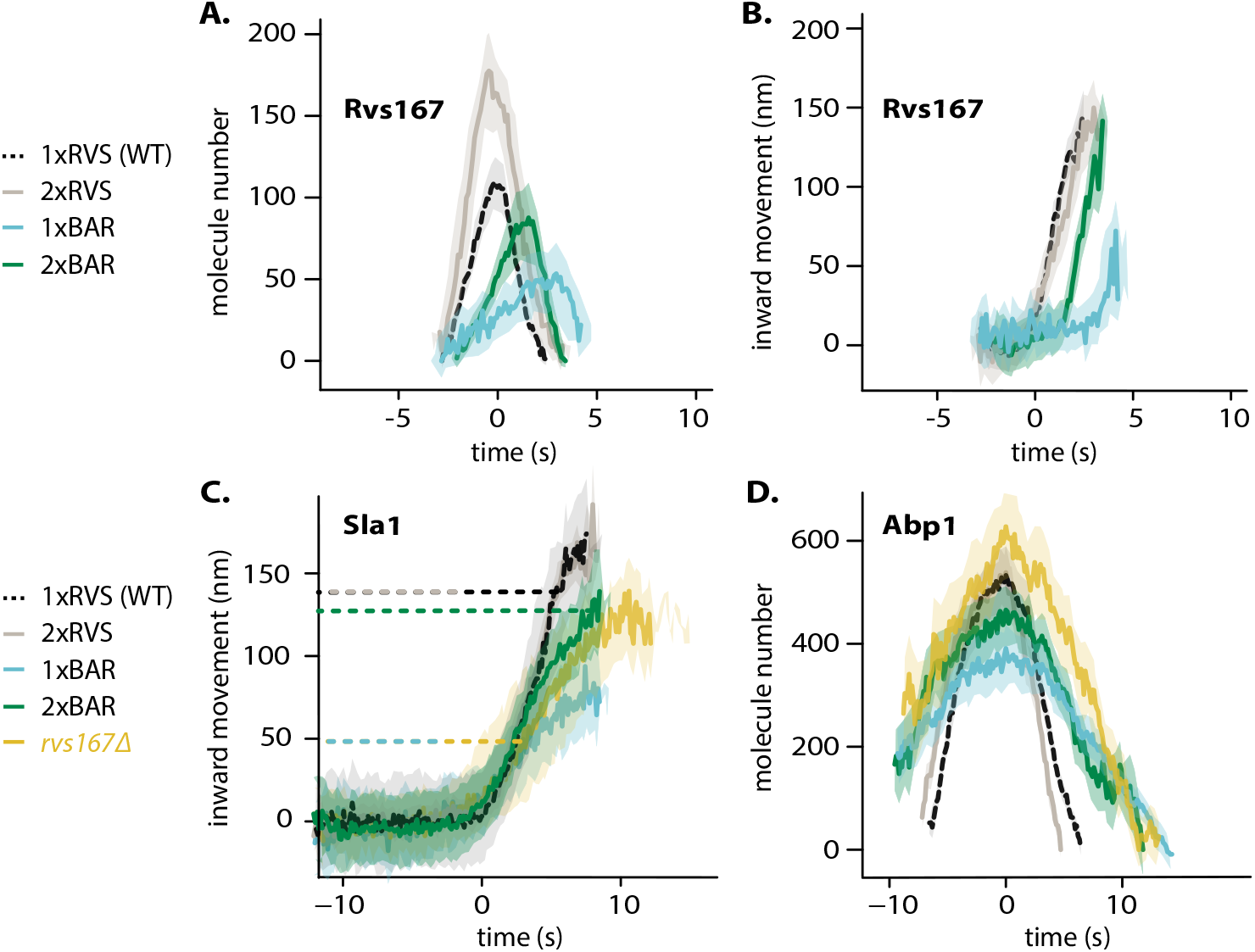
Titration of BAR molecules. **A,B:** Molecule numbers and centroid positions of Rvs167. Centroid positions were co-aligned with Abp1-mCherry so that time=0 s corresponds to Abp1 intensity maximum. Centroids movements were aligned in the y-axis to a starting position = 0 nm. **C:** Sla1 centroid positions, aligned so that the centroids begin inwards movement at the same time. Aligned in the y-axis so that y=0 nm corresponds to non-motile centroid position. Dashed lines correspond to the Sla1 centroid positions when intensity of Abp1 in the corresponding strain is at maximum. **D:** Abp1 molecule number, aligned that time=0 s corresponds to Abp1 intensity maximum

**Figure 7.**
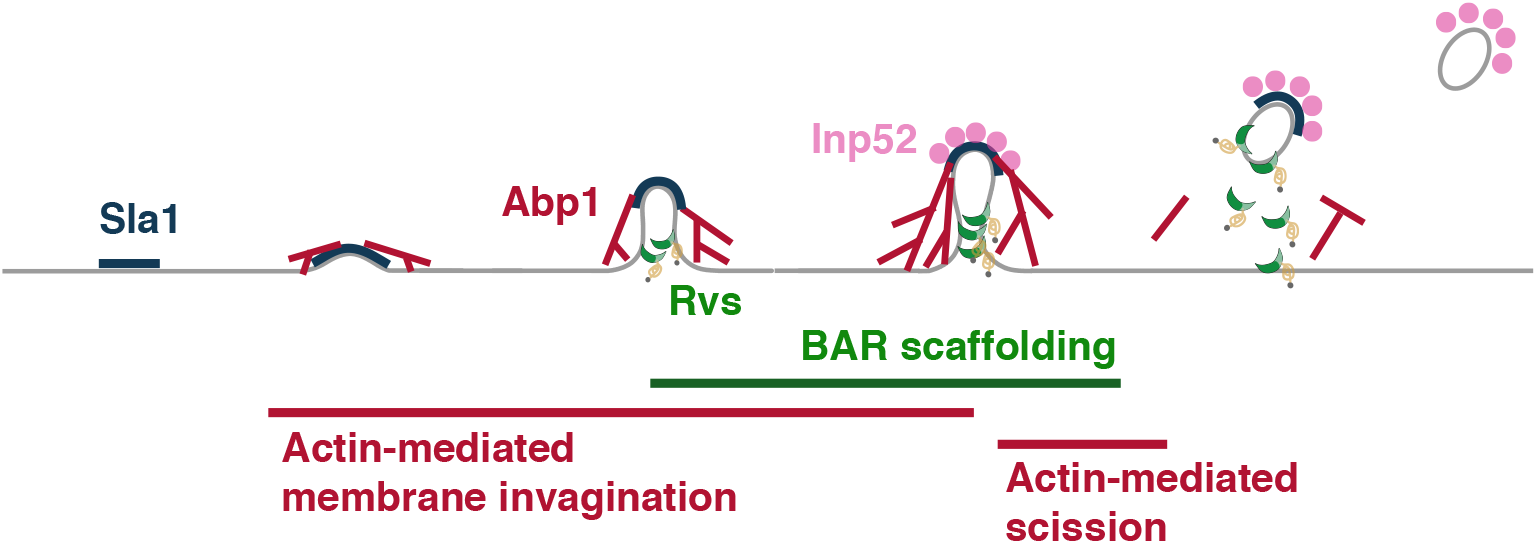
Model for yeast endocytic scission. Membrane at an endocytic site is bent by forces derived from actin polymerization. BAR domains arrive at a tubular invagination and scaffold the membrane, delaying scission. Actin forces eventually overcome the influence of BAR scaffolding, and the membrane breaks, resulting in vesicle formation.

Maximum number of WT and mutant Rvs167 molecules recruited at endocytic sites varied in 1xRVS (WT), 2xRVS, 1xBAR, 2xBAR strains between 50 and 180 copies (Fig.6A, S6). Excess Rvs recruited in 2xRVS cells (compared to 1xRVS) did not change the total movement of Rvs167, but Rvs disassembly took longer (Fig.6A). In the 2xBAR case, amount of Rvs167-sh3Δ molecules recruited to endocytic sites increased compared to 1xBAR, as did the movement of the centroid (Fig.6A, B). BAR domain recruitment increased from 1xBAR, to 2xBAR, 1xRVS, and was finally maximal in 2xRVS cells. The trend of inward movement of the Rvs167 centroid suggests that movement correlates with number of BAR molecules recruited to sites, but saturates in the case of 2xRVS. The delayed disassembly of 2xRVS compared to 1xRVS may be due to a change in interaction between the BAR domains and underlying membrane. The membrane may be already saturated with bound RVS, causing perhaps interaction between Rvs dimers rather than Rvs and membrane. Alternatively, excess Rvs molecules may be recruited to the vesicle, so there is a delay in disassembly of these molecules compared to those on the invagination tube.

Abp1 molecule numbers and lifetimes at endocytic sites were different between the 1xRVS 2xRVS, 1xBAR, 2xBAR strains (Fig.6D, S5). Total Abp1 molecules recruited were reduced in 1xBAR compared to the 2xBAR, 1xRVS and 2xRVS (Fig.6D, Fig.S6). As Abp1 molecule numbers increased, shorter lifetimes, approaching that of WT Abp1 were observed (Fig.S5). Comparing between the 1xRVS (WT), 2xRVS, 1xBAR, 2xBAR strains, the cells with higher Abp1 molecule numbers showed larger total movement of the Sla1 centroid (Fig.6C,D). This indicates a correlation between the maximum number of Abp1 molecules recruited and total invagination length. In *rvs167*Δ cells, measured Abp1 molecule numbers were about the same as in WT (Fig.6D). Quantification of Abp1 molecule numbers in these cells is confounded by the existence of two types of endocytic events: successful and retracting events. We were unable to separate these events in the molecule number quantification, but we speculate that retracting events may continue to assemble an actin network after or during retraction. This accumulation could compensate for smaller Abp1 numbers that may have been measured at successful endocytic events. A much smaller percentage of endocytic events in 1BAR and 2xBAR retract, so we do not expect retracting endocytic invaginations in these cells to confound the Abp1 quantification significantly (Fig.S6).

We found that increasing the expression of RVS caused increased recruitment of Rvs molecules to endocytic sites. We also found that the number of Rvs BAR domains recruited to membrane invaginations correlates positively with total length of the invagination. Furthermore, this length of the invagination also correlates positively with the number of Abp1 molecules recruited.

## Discussion

Recruitment and function of the Rvs complex has been studied in this work, and the applicability of several membrane scission models to yeast endocytosis have been tested. We propose that Rvs is recruited to endocytic sites via interactions between the BAR domains and invaginated membrane, and also via SH3 mediated protein-protein interactions. SH3 interactions are required for efficient recruitment of Rvs. We found that assembly of Rvs at the membrane invagination delays membrane scission, allowing the invagination to grow to its full length. WT invagination growth depends on recruitment of a critical number of Rvs molecules. Both timing and recruitment efficiency of Rvs appear crucial to Rvs function.

### BAR domains sense in vivo membrane curvature and time the recruitment of Rvs

The curved structure of Endophilin and Amphiphysin BAR domains allows them to interact with curved membranes. These proteins are able to form organized assemblies on tubular membranes in vitro (***Mim et al., 2012***). Rvs167-sh3Δ localized to endocytic sites when curvature was present (Fig.3B, “Untreated”). Without the SH3 domain, and in the absence of membrane curvature in *sla2*Δ cells, Rvs167-sh3Δ did not localize to endocytic sites (Fig.3B, “*sla2*Δ”). This indicates that the Rvs167-sh3Δ localization was via BAR-membrane curvature interaction. This demonstrates that the Rvs BAR domain senses and requires membrane curvature to interact with endocytic sites. Rvs167-sh3Δ had a similar average lifetime at endocytic sites as full-length Rvs167 (Fig.5A,B). However, time alignment with Abp1 showed that there was a delay in the recruitment of Rvs167-sh3Δ (Fig.5C,D). Sla1 centroid movement was slower in *rvs167-sh3*Δ cells than in WT cells: it takes longer for the membrane in these cells to reach the same invagination length as WT. We propose that the start of Rvs recruitment is timed to a specific membrane invagination length - therefore to a specific membrane curvature - accounting for the delay in recruitment of Rvs167-sh3Δ. The precise timing of recruitment is therefore provided by the BAR domain interacting with membrane at a specific curvature.

### SH3 domains allow efficient, actin- and membrane curvature-independent recruitment of Rvs

Rvs167-sh3Δ accumulated to about half the number of full length Rvs167 (Fig.5C,D) even though similar cytoplasmic concentration was measured for both proteins (Fig.S4), indicating that loss of the SH3 domain decreases the efficiency of recruitment of Rvs to endocytic sites. In *sla2*Δ cells, full-length Rvs167 forms patches on the membrane (Fig.3B, “*sla2*Δ”). Since Rvs167-sh3Δ does not localize to the plasma membrane in *sla2*Δ cells, localization of the full-length protein must be mediated by the SH3 domain. The full-length Rvs167 is able to assemble and disassemble at cortical patches in *sla2*Δ cells, that is, without the curvature-dependent interaction of the BAR domain (Fig.S3). This indicates recruitment and disassembly of Rvs can occur via interactions between its SH3 domains and endocytic sites. In *sla2*Δ cells treated with LatA (Fig.3B, “*sla2*Δ+ LatA”), both membrane curvature and actin are removed from endocytic sites. Full-length Rvs167 in these cells still shows transient localizations at the plasma membrane. Therefore the SH3 domain is able to localise the Rvs complex in an actin- and curvature-independent manner.

### Recruitment of Rvs167 affects endocytic actin network dynamics

In WT cells, the Abp1 and Rvs167 fluorescent intensities peaked concomitantly (Fig.5C,D), and the consequent decay of both coincided. Membrane scission occurs around the intensity peak of Rvs167 (***Kukulski et al., 2012***; ***Picco et al., 2015***). Coincident disassembly therefore indicates that upon vesicle scission, the actin network is rapidly disassembled. This coincident peak was lost in *rvs167-sh3*Δ cells: Rvs167-sh3Δ fluorescent intensity peaks after Abp1 intensity starts to drop. The decay of Abp1 is also prolonged, taking over double the time as in WT. Although it is not clear what the decoupling of Abp1 and Rvs167-sh3Δ peaks means, the changes in Abp1 dynamics suggests a strong disruption of the actin network. In 1xBAR cells, the average lifetime of actin marker Abp1 was about 25 s (Fig.S5). This lifetime decreases in 2xBAR cells to about 20 s, a shift towards the WT Abp1 lifetime of around 10 s. Therefore we conclude that recruitment of the Rvs BAR domains to the invagination regulates actin network dynamics.

### Rvs acts as a membrane scaffold, delaying membrane scission

Invagination length in non-retracting endocytic events in *rvs167*Δ cells at scission time was about 50 nm (Fig.1E), only a third the WT length. Together with electron microscopy data (***Kukulski et al., 2012***), this shows that scission can occur at much shorter invagination lengths. In WT cells, scission does not occur at these lengths, instead invaginations grow to 150 nm (***Kukulski et al., 2012***). Since invagination lengths were increased, compared to *rvs167*Δ and 1xBAR, by overexpression of the Rvs167-sh3Δ protein, that is, in 2xBAR (Fig.6A,C), we think that localization of Rvs-BAR domains to the membrane tube stabilizes the membrane and allows invaginations to progress (***Boucrot et al., 2012***; ***Dmitrieff and Nédélec, 2015***). Yeast endocytosis is heavily dependent on a dynamic actin network to generate the forces that bend the membrane (***Kübler et al., 1993***; ***Kaksonen et al., 2003***; ***Picco et al., 2018***). We propose that Rvs accumulation stabilizes the membrane invagination and thereby also increases the amount of actin required to sever the membrane. This allows the invagination to grow until WT invagination length is reached. We speculate that continued invagination growth allows the actin network to generate enough force to compensate for the stabilization. There is a limit to the stabilization by BAR domains: in 2xRVS cells, invagination lengths are the same as in 1xRVS cells even though more Rvs is recruited. It is possible that the nature of interaction of the Rvs complex with the membrane changes after a certain amount of Rvs is recruited. Once the membrane is saturated with Rvs molecules, BAR domains may interact with each other rather than with the underlying membrane. This could explain the changes in the disassembly dynamics of Rvs in the 2xRVS case (Fig.6A).

If enough forces are generated at around 50 nm, why is scission inefficient and membrane retraction rates increased in *rvs167*Δ compared to WT? Forces generated by the actin network may be at a threshold level when the invaginations are short. There could be enough force to sever the membrane, but not enough to sever reliably. The Rvs scaffold may then stabilize the membrane invagination, preventing retraction, and allowing continued growth. This subsequently allows the actin network to continue growing, accumulating actin. Eventually enough actin is accumulated to reliably cause scission. We hypothesise that increased actin amount yields higher force on the membrane. This force stretches the membrane, eventually breaking it. Controlling membrane tube length could also be a way for the cell to control the size of the vesicles formed, and therefore the amount of cargo that can be packed into the vesicle.

### What causes membrane scission?

We have tested candidate proteins implicated in yeast endocytic scission and looked for scission defects. Increased Sla1 retraction rates would indicate higher rate of scission failure. Larger total movement of Sla1 and Rvs167 centroids would indicate that a longer invagination has been formed, and that scission has not occured at normal invagination lengths. We did not see a change in Sla1 or Rvs167 centroid movements that would indicate scission defects in any mutants that we studied, other than in Rvs mutants. In *vps1*Δ cells, there is no major change in retraction rate, nor are there changes in Sla1 or Rvs167 dynamics. We conclude that Vps1 is not necessary for Rvs localization or function, and is not necessary for scission.

Sla1 and Rvs167 centroid dynamics showed that deletion of neither Inp51 nor Inp52 resulted in scission delay. In *inp51*Δ cells, Rvs167 assembly and disassembly was slightly slower than in WT: Inp51 could play a role in recruitment to and release of Rvs from endocytic sites. In the *inp52*Δ cells, about 12% of Sla1-GFP tracks retracted. Inp52 has a moderate influence on scission efficiency, but this is not reflected in our observation of invagination dynamics. In *inp52*Δ cells, Sla1 assembly is slower than in WT, and Sla1 and Rvs167 centroids persisted after scission. Inp52 likely plays a role in assembly of coat proteins, and in recycling endocytic proteins from the vesicle to the cytosolic pool. Synaptojanins could help recruit Rvs at endocytic sites via their proline-rich domains by binding Rvs167 SH3 domains. They are involved in vesicle uncoating post-scission, likely by dephosphorylating PI(4,5)P_2_ and inducing disassembly of PI(4,5)P_2_-binding endocytic proteins. The synaptojanins do not appear to play a major role in scission, but Inp51 and Inp52 may function synergistically to influence membrane tension. The compounded problems related to lipid hydrolysis, and lack of tools that have the time resolution to measure membrane tension in vivo prevent us from conclusively ruling out line tension as a contributor to yeast endocytic scission.

Our RVS duplication data is able to test whether the protein friction model is applicable to yeast endocytic scission (***Simunovic et al., 2017***). According to this model, a frictional force between a moving membrane tube and a coat of BAR protein bound to it causes the tube to undergo scission. Therefore, a higher frictional force should break the tube sooner than a lower force. We increased the frictional force on the membrane by increasing the number of BAR domains bound to the membrane tube, in 2xRVS cells. In 2xRVS cells, adding up to 1.6x the WT amount of Rvs at faster rates to membrane tubes did not affect the length at which the membrane undergoes scission (Fig.6). In *rvs167*Δ cells, frictional forces generated should be reduced compared to WT cells. Rather than increased Sla1 movement as this model would predict, we observed decreased Sla1 movement (Fig.1). We therefore think that protein friction does not contribute significantly to membrane scission in yeast endocytosis.

A similar amount of Abp1 is recruited in both 1xRVS and 2xRVS cases, corresponding to coat movement of about 150 nm. Magnitude of coat movement correlates with the total amount of Abp1, and therefore, with the amount of actin recruited. A dynamic actin network is required for endocytosis in yeast (***Kübler et al., 1993***; ***Picco et al., 2018***), and such a network is able to generate force (***Theriot et al., 1992***). Coupling between the actin network and membrane is necessary for invagination formation (***Skruzny et al., 2012***; ***Picco et al., 2018***). The current understanding of yeast endocytosis suggests that the membrane is pushed into the cytoplasm by an actin network polymerizing at the base of the invagination, and is mechanically coupled to the invagination tip. More actin recruitment can generate higher force (***Bieling et al., 2016***). Actin may also provide a scaffold that aids membrane invagination. More actin is therefore consistent with scaffolding as well as with increased force generation. We propose that increased Abp1 recruitment - and therefore increased actin - leads to an increasing pushing force on the membrane, and that this force is responsible for invagination growth as well as for membrane scission. Stretching the membrane can eventually cause it to break, causing vesicle formation. The amount of force necessary to break the membrane is determined by properties of the membrane like rigidity and tension, properties of the proteins accumulated on the membrane, and by the high intracellular turgor pressure in yeast cells (***Dmitrieff and Nédélec, 2015***). Once this force is overcome, vesicle scission occurs, and membrane-bound Rvs is released. On the other hand, release of Rvs could cause instabilities in membrane shape that could also lead to scission (***Dmitrieff and Nédélec, 2015***). It is unclear whether scission causes release of Rvs, or vice-versa. The observation that Rvs167 can accumulate and disassemble on the membrane in the absence of membrane curvature (in *sla2*Δ cells) suggests that binding-unbinding can be mediated by another interaction partner. This in turn can allow speculation that Rvs release can be triggered by this partner. A method for detecting scission with high temporal resolution is needed to resolve whether Rvs release or scission occurs first. Release of the SH3 domains could eventually indicate to the actin network that vesicle scission has occurred, influencing disassembly of actin components.

We propose that Rvs is recruited to endocytic sites by two distinct mechanisms. The SH3 domain of Rvs167 recruits Rvs to endocytic sites, effectively increasing the likelihood of BAR domain interaction with tubular membrane. BAR domains bind endocytic sites by sensing tubular membrane. The membrane invagination is stabilized against scission by BAR-membrane interaction. This stabilization prevents actin-generated forces from rupturing the membrane, and the invaginations continue to grow in length as actin continues to polymerize. As actin continues to accumulate, pushing forces overcome the resistance to membrane scission. The membrane ruptures, and a vesicle is formed.

## Methods and Materials

### Homologous recombination with PCR casette insertion

Tagging or deletion of endogenous genes was done by homologous integration of the product of a Polymerase Chain Reaction (PCR) using appropriate primers and a plasmid containing a selection cassette and fluorescent tag, or only selection cassette for gene deletions (***Janke et al., 2004***). PCRs used the Velocity Polymerase for fluorescent tagging, and Q5 for gene deletions using the NAT casette. All fluorescently tagged genes have a C-terminus tag and are expressed endogenously. Gene deletions and fluorescent tags are checked by PCR. *vps1*Δ and gene duplications were confirmed by DNA sequencing.

### Live-cell imaging

#### Sample preparation for live imaging

Yeast cells were grown overnight at 25°C in imaging medium Synthetic Complete without L-Tryptophan (SC-Trp). 40 μL 4mg/ml Concavalin A (ConA) was incubated on a coverslip for 10 minutes. 40 μL yeast cells at OD_600_= 0.3-0.8 was added to the coverslip after removing the ConA, and incubated for another 10 minutes. Cells were then removed, adhered cells were washed 3x in SC-Trp, and 40 μL SC-TRP was finally added to the coverslip to prevent cells from drying.

#### Sample preparation for live imaging in LatA treated cells

Cells went through the same procedure as above till the last washing step. Instead of SC-Trp, 100x diluted LatA in SC-Trp was added to the adhered cells. Cells were incubated in LatA for 10 minutes before imaging.

#### Epifluorescent imaging for centroid tracking

Live-cell imaging was performed as in our previous work (***Picco et al., 2015***). All images were obtained at room temperature using an Olympus IX81 microscope equipped with a 100×/NA 1.45 PlanApo objective, with an additional 1.6x magnification lens and an EMCCD camera. The GFP channel was imaged using a 470/22 nm band-pass excitation filter and a 520/35 nm band-pass emission filter. mCherry epifluorescence imaging was carried out using a 556/20 nm band-pass excitation filter and a 624/40 band-pass emission filter. GFP was excited using a 488 nm solid state laser and mCherry was excited using a 561 nm solid state laser. Hardware was controlled using Metamorph software. For single-channel images, 80-120 ms was used as exposure time. All dual-channel images were acquired using 250 ms exposure time. Dual-color images were obtained using simultaneous illumination using 488 and 561 nm lasers. A dichroic mirror and emission filters 650/75 and 525/50 were used for image acquisition, and corrected using TetraSpeck beads for chromatic abberation.

#### Epifluorescent imaging for molecule number quantification

Images were acquired as in previous work (***Picco et al., 2015***). Z-stacks of cells containing the GFP- or mCherry-tagged protein of interest, incubated along with cells containing Nuf2-GFP, were acquired using 400 ms exposure using a mercury vapour lamp, on a CCD camera. Z stacks were spaced at 200 nm.

### Live-cell image analysis

Images were processed for background noise using a rolling ball radius of 90 pixels. Particle detection, and tracking was performed for a particle size of 6 pixels, using scripts that combine background subtraction with Particle Tracker and Detector, that can be found on ImageJ (http://imagej.nih.gov). Further analysis for centroid averaging, alignments between dual-color images and single channel images, for alignment to the reference Abp1 were done using scripts written in Matlab (Mathworks) and R (www.r-project.org), written originally by Andrea Picco, and modified by me. Details of analysis can be found in previous work (***Picco et al., 2015***). All movement and intensity plots from centroid tracking show the average centroid with 95% confidence interval.

### Quantification of cytoplasmic concentration

On a maximum intensity projection of time-lapse images, the average pixel intensity within a circle of set radius in the cytoplasm was measured. This circle is manually arranged so that cortical patches were excluded, and mean intensity was acquired for about 10 cells of each cell type. A fixed area outside the cells was drawn, and mean intensity was calculated to establish “background intensity”. This background intensity was then subtracted from the mean intensity to obtain a rough measure of cytoplasmic intensity.

## Acknowledgments

We would like to thank the entire Kaksonen lab, especially Daniel Hummel, Andrea Picco, and Mateusz Kozak for critical reading of this manuscript. This work was supported by the Swiss National Science Foundation (grant 310030B_182825) and by the NCCR Chemical Biology funded by the SNSF.

## Supplementary Material

**Figure S1.**
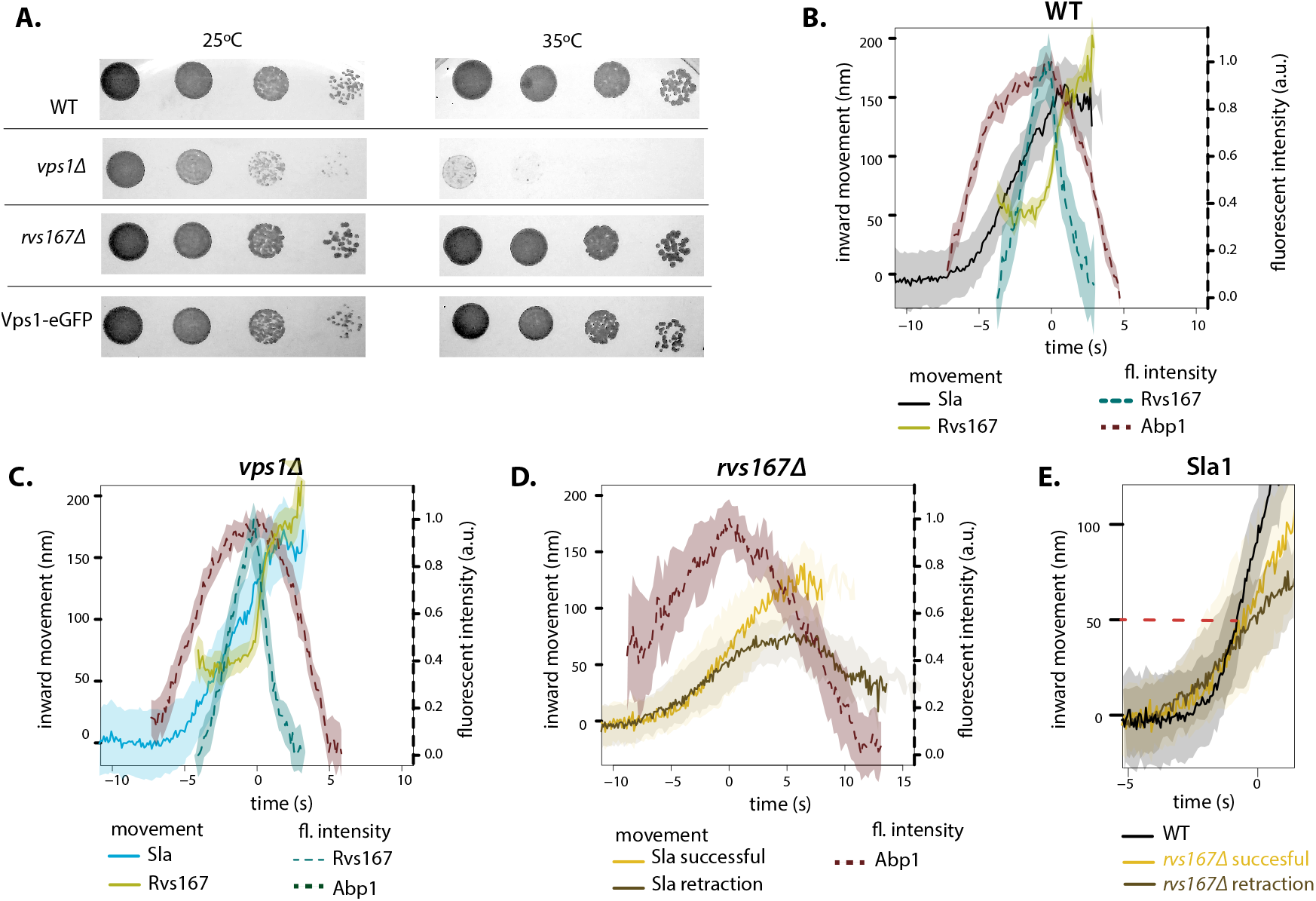
A: Growth assay of WT, *vps1*Δ, *rvs167*Δ, and cells expressing Vps1-eGFP at 25 °C and 35 °C. B, C, D: Sla1 and Rvs167 centroids aligned so that time=0 s is the maximum of Abp1 fluroescent intensity. Centroid movements aligned so that y=0 nm is the starting Sla1 position. Normalized Abp1 and Rvs167 fluorescent intensities in WT, *vps1*Δ, and *rvs167*Δ cells. E: Sla1 centroids in WT, and successful and retracting Sla1 in *rvs167*Δ cells. Centroids are aligned so that WT Sla1 begins inwards movement at the same time as *rvs167*Δ Sla1. Dashed line indicates 50 nm.

**Figure S2.**
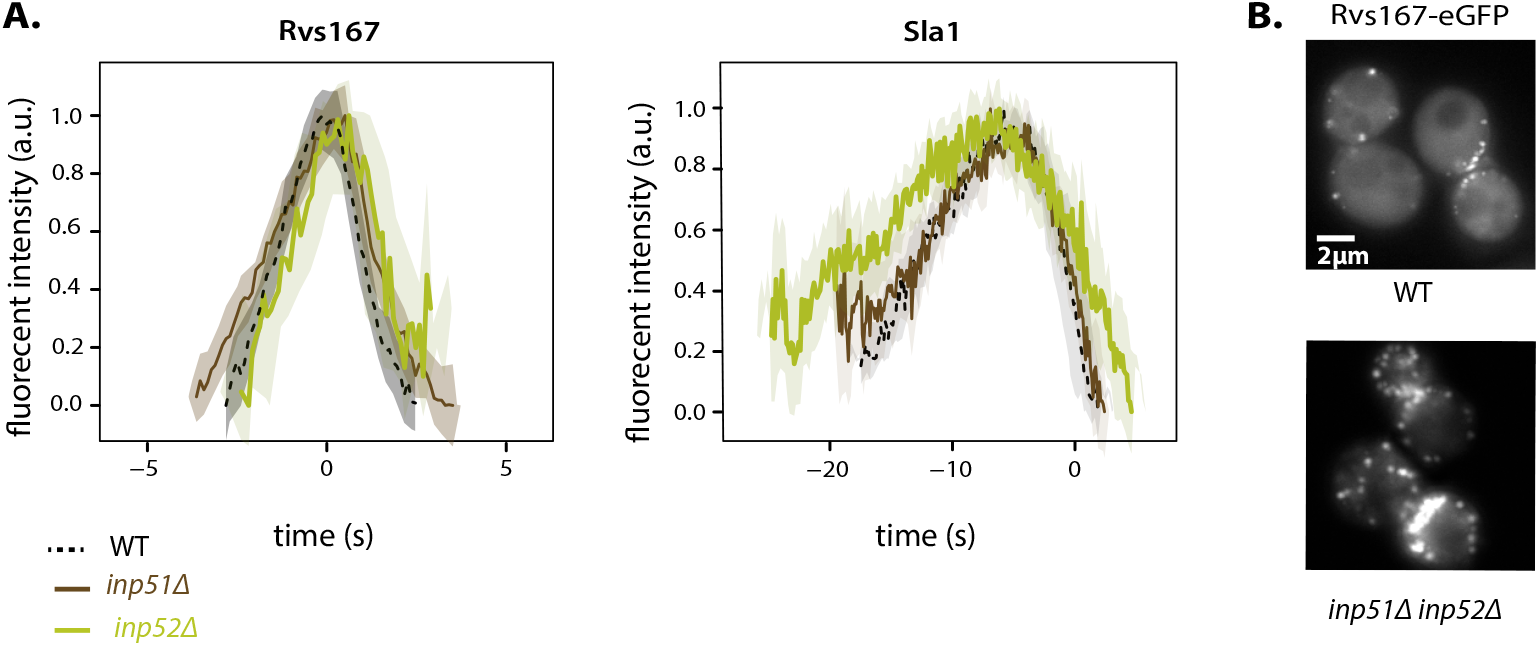
A: Normalized Rvs167 and Sla1 fluorescent intensities in synaptojanin deletion aligned in time so that time=0 s corresponds to Abp1 intensity maximum. B: Maximum intensity projection of a time-lapse movie of Rvs167-eGFP in WT and *inp51*Δ*inp52*Δ *cells.*

**Figure S3.**
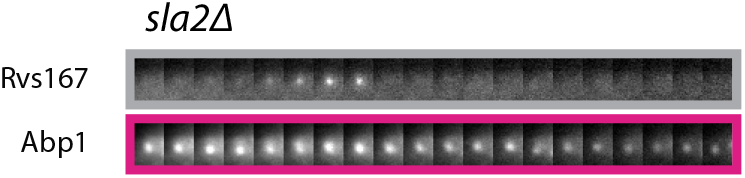
Rvs167-eGFP and Abp1-mCherry in *sla2*Δ cells.

**Figure S4.**
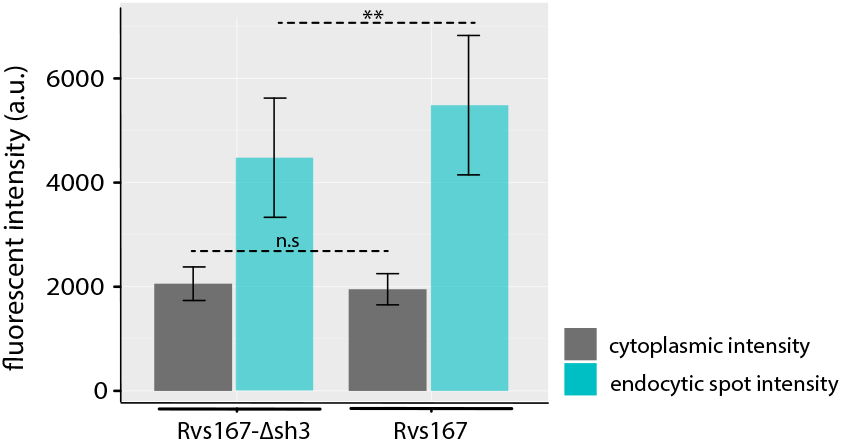
Cytoplasmic intensity and intensity at endocytic patches of Rvs167-eGFP and Rvs167-sh3 Δ-eGFP. Error bars are standard deviation, p values from two-sided t-test, *= p ≤0.05, **= p≤0.01, ***=p ≤0.001.

**Figure S5.**
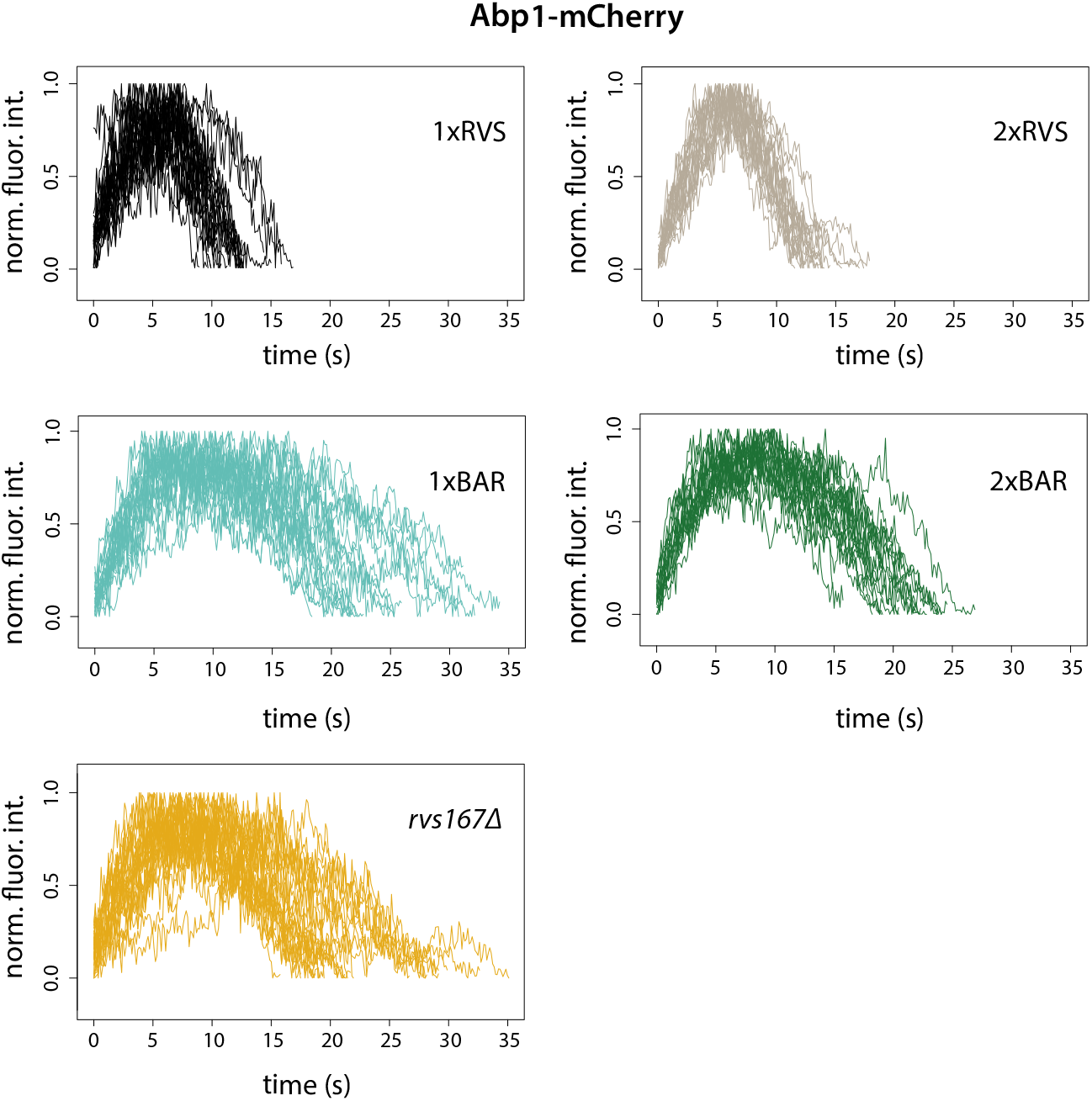
Normalized fluorescent intensities of Abp1-mCherry in 1xRVS, 2xRVS, 1xBAR, 2xBAR and *rvs167*Δ cells.

**Figure S6.**
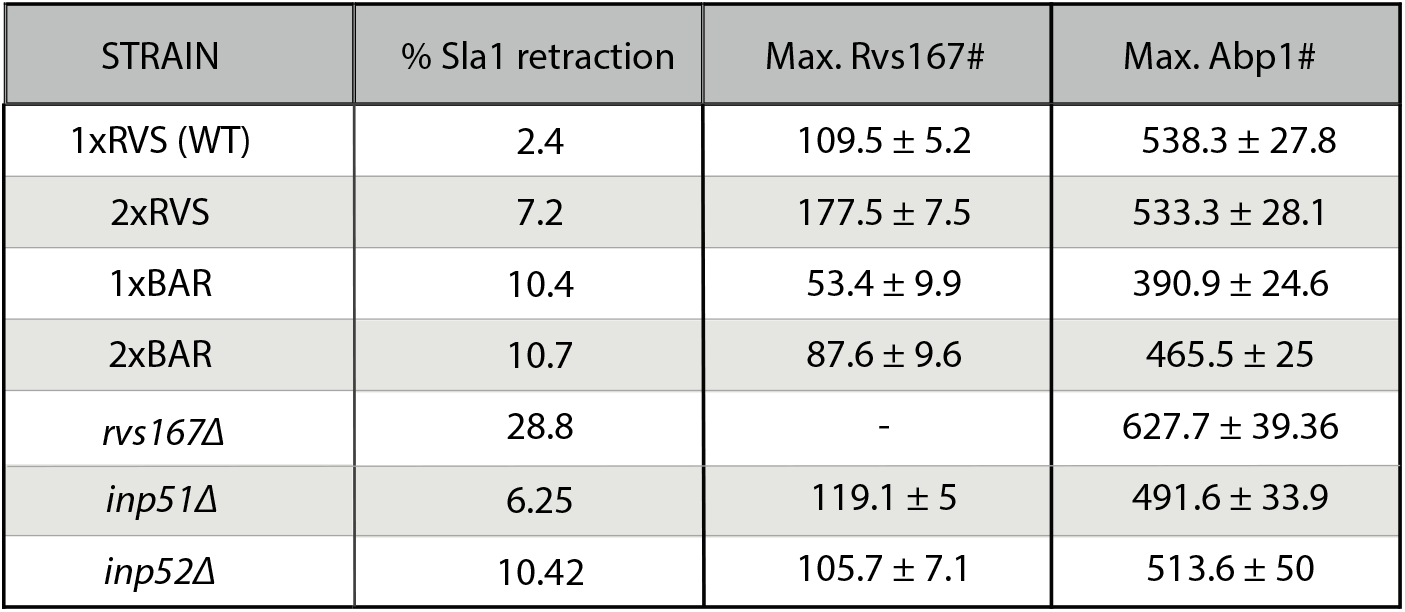
Percentage of retracting events, Rvs167 and Abp1 molecule numbers

